# Population dynamics and age-dependent mortality processes in tropical reptiles

**DOI:** 10.1101/575977

**Authors:** Hugo Cayuela, Godfrey C. Akani, Emmanuel M. Hema, Edem A. Eniang, Nioking Amadi, Stephanie N. Ajong, Daniele Dendi, Fabio Petrozzi, Luca Luiselli

**Affiliations:** Institut de Biologie Intégrative et des Systèmes (IBIS), Université Laval, Québec, QC G1V 0A6, Canada; Department of Applied and Environmental Biology, Rivers State University of Science and Technology, P.M.B. 5080, Port Harcourt, Rivers State, Nigeria; Institute for Development, Ecology, Conservation and Cooperation, via G. Tomasi di Lampedusa 33, I-00144 Rome, Italy; Université de Dédougou, UFR/Sciences Appliquées et Technologiques, Dédougou, Burkina Faso; Laboratoire de Biologie et Ecologie Animales, Université Ouaga I Prof. Joseph Ki-Zerbo, Ouagadougou, Burkina Faso; Department of Forestry and Wildlife, University of Uyo, Uyo, Akwa Ibom State, Nigeria; Department of Fisheries, Lagos State University, Lagos, Nigeria; Department of Zoology, University of Lomé, Lomé, Togo; Istituto Tecnico di Ecologia Applicata, Fano (PU), Italy

**Keywords:** age-dependent mortality, senescence, survival, recruitment, reptile

## Abstract

Understanding age-dependent mortality processes is a critical challenge for population biologists. Actuarial senescence appears to be a common process across the tree of life. Senescence patterns are highly variable in pluricellular organisms: senescence can be gradual or sharp and its onset may be early or delayed. By contrast, studies revealed that organisms may also not experience senescence at all while others display a “negative senescence”; i.e. a decrease of mortality rate with age. To date, studies on senescence have largely focused on human and other endotherm vertebrates, limiting our understanding of senescence in amniotes as a whole. By contrast, few have examined the diversity of senescence patterns in ectotherm vertebrates as reptiles. Here, we examined population dynamics and age-dependent mortality patterns in three tropical tortoises (*Kinixys erosa, Kinixys homeana, Kinixys nogueyi*) and snakes (*Bitis gabonica, Bitis nasicornis, Causus maculatus*). Our study revealed that tortoises of *Kinixys* genus had a higher survival and a lower recruitment than snakes of the genera *Bitis* and *Causus*, indicating that they have a slower life history. Furthermore, we showed that survival more slowly decreased with age in tortoises than in snakes. In addition, we highlighted contrasted patterns of age-dependent mortality in the three genera. In *Kinixys*, the relationship between mortality rate and age was positive and linear, suggesting gradual senescence over tortoise lifetime. By contrast, the relationship between mortality rate and age was negative and sharp in *Bitis* and *Causus*, possibly due to a “negative senescence” starting early in life. Our study highlighted various age-dependent mortality patterns in tropical reptiles. It also contributed to extend our knowledge of senescence in ectotherm vertebrates whose the demography is still poorly understood. In addition, while negative senescence has never been reported in endotherm vertebrates, our results showed that it can be common phenomenon in ectotherms.

## Introduction

The ageing theories state that an organism’s survival should decrease with age, a phenomenon called actuarial senescence (hereafter senescence) (Hamilton 1966, Monaghan et al. 2008). In the 1950s, Medawar introduced the mutation accumulation theory that predicts that the strength of natural selection decreases with age after the primiparity. It states that the efficiency of the purging of deleterious mutations – having a detrimental effect on fitness components including survival – diminishes with age. Simultaneously, Williams (1957) proposed the theory of antagonistic pleiotropy postulating that senescence is a by-product of selection: an allelic variant conferring a selective advantage at early stage may lead to a decreased survival later in life. In the 1970s, Kirkwood (1977) introduced the theory of disposable soma that poses that senescence results from a trade-off between an early reproduction and somatic maintenance. A decrease in the energy allocation in somatic maintenance for the benefit of reproduction leads to lower survival-related performances and senescence.

More recently, studies suggested that senescence is a common process across the tree of life (Baudisch et al. 2013, Jones et al. 2014, Colchero et al. 2019; but see Jones & Vaupel 2017). Senescence patterns are highly variable in pluricellular organisms: it can be gradual or sharp and its onset may be early or delayed (Jones et al. 2014, Colchero et al. 2019). Studies showed that the onset of senescence is usually associated to species position along the fast-slow continuum of life histories (Stearns 1992, Oli 2004, Bielby et al. 2007). Species at the slow end of the continuum – with a long lifespan, a low fecundity, and a delayed maturity – usually have a delayed senescence (Jones et al. 2008, Kiørboe et al. 2015, Salguero-Gómez & Jones 2017). However, several species do not experience senescence at all (Jones et al. 2014, Jones & Vaupel 2017, Colchero et al. 2019). Moreover, other organisms may have a “negative senescence” (Vaupel et al. 2004, Jones & Vaupel 2017), a phenomenon that usually occurs in species exhibiting large body size variation over life and size-dependent survival. Mortality increases as body size decreases and size increases with age, which results in a negative relationship between mortality and age (see several cases in Jones et al. 2014).

To date, studies dealing with age-dependent mortality processes and senescence in amniotes have broadly focused on human and other endotherm vertebrates (Nussey et al. 2013, Fridlyanskaya et al. 2015, Shefferson et al. 2017). By contrast, the senescence patterns of ectotherm vertebrates as reptiles have been overlooked for a long time (Robert & Bronikowski 2010, Colchero et al. 2019), thus limiting our understanding of senescence processes to a restricted set of amniotes. The reptile class is the second most species-rich group of amniotes after birds and hosts 32% of the tetrapod diversity (Pincheira-Donoso et al. 2012, IUCN 2019). To date, few studies suggested that senescence patterns could be highly variable in reptiles: three of them indicated that senescence may occur or not in squamates and turtles (Robert & Bronikowski 2010, Warner et al. 2016, Colchero et al. 2019) while another one showed that negative senescence can be found in tortoises (Jones et al. 2014). Yet, the small number of studies focusing on this topic is insufficient to reflect the potentially high diversity of senescence patterns in reptiles.

Here, we examined population dynamics and age-dependent mortality patterns in three tropical testudinid tortoises (*Kinixys erosa, Kinixys homeana, Kinixys nogueyi*) and viperid snakes (*Bitis gabonica, Bitis nasicornis, Causus maculatus*). The six species were surveyed using capture-recapture method over a 16-years period in a tropical forest of western Africa (Nigeria). First, we quantified adult survival and recruitment in the six species of tortoises and snakes. Based on previous studies on the demography of testudines (Gibbons 1987; Congdon et al. 1993, 1994), we expected (1) tortoises to have slow life histories (i.e. higher adult survival and lower recruitment). By contrast, we did not have any precise expectations for *Bitis* and *Causus* snakes (their demographic parameters are unknown as in most of tropical snakes), even if the lifespan measured in captivity suggests a long lifespan for *Bitis* species. Second, we examined age-dependent survival and mortality rate. After showing that tortoises have slower life histories than snakes, we hypothesized that (2) survival should decrease more slowly with age in tortoises than in snakes. We also hypothesized that (3) if senescence occurs in both tortoises and snakes, tortoises should have a more delayed senescence.

## Materials and methods

### Studied species

*Kinixys homeana, K. erosa* and *K. nogueyi* are omnivorous tortoises inhabiting the forested areas and the forest-plantation mosaics of West Africa. They feed essentially on mushrooms and invertebrates, and are rapidly declining because of habitat loss and overharvesting for local consumption (Luiselli & Diagne 2013, 2014). They occur in sympatry in several forest zones of the Niger Delta and of the Togo hills (Luiselli & Diagne 2013, 2014).

*Bitis gabonica* and *Bitis nasicornis* are two massive viper species (usually longer than 130 cm), with a wide distribution across the Guinea-Congolian forest belt, where they inhabit forest and forest-plantation mosaics (Chippaux 2013). Their diet is based essentially on rodents, and, in southern Nigeria, it is very similar in sympatric conditions (Luiselli & Akani 2003). *Causus maculatus* is a small viper species (up to 60 cm in length), nocturnal in habits, that feed mainly of frogs and inhabit forest patches as well as highly disturbed areas and plantations in West Africa (Chippaux, 2013).

### Study area and capture-recapture surveys

The field study was performed in the Port Harcourt area of the Niger Delta, Rivers State, south-eastern Nigeria. The study area is heavily populated with hundreds of villages interspersed by patches of forests and cultivated lands (yam, cassava and pineapples). The climate of the study region is tropical, with well-delineated dry (from November to March) and wet (from April to October) seasons. Mean annual rainfall averages around 4000 mm, making it one of the wettest areas in Africa. The wet season peaks in July, and the dry season peaks in January and February. Relative humidity rarely dips below 60% and fluctuates between 90% and 100% for most of the year. During most of the rainy season cloud cover is nearly continuous, with about 1500 mean annual sunshine hours and an average annual temperature of approximately 28°C. Both vipers and tortoises are particularly active above-ground by wet season whereas they spend most of the dry season months hidden, with only nocturnal activity on occasion (Luiselli 2003a, 2006a).

The survey was conducted between 2000 and 2016: tortoises were surveyed over the complete period while snakes were monitored from 2000 to 2007. M-array matrices documentig the mark-recapture process in the six species is provided in Supplementary material, Table S1-S4. Most of the surveys were done between 0630-1030 hour and at 1730-2230 hour (Lagos standard time). Field research was suspended during the central daylight hours because of too much hot, and therefore no activity above-ground of these reptiles. Snakes and tortoises were studied simultaneously, as they were sympatric and syntopic inside the same forest patch. These reptiles were searched for by means of different surveying procedures: (i) random searching along all appropriate forest micro-habitats, (ii) pitfalls with drift fences checked every day, and (iii) examination of specimens just captured by local people that were employed by us to help in getting more individuals from the field. Overall, random searching was done during 911 different days, 518 during the wet season (May to September), and 393 during the dry season (October to April). Every tortoise was identified to species, sexed and individually marked by unique sequences of notches filed into the marginal scutes. Each snake individual was permanently marked by ventral scale clipping. Tortoises were generally easier to locate than vipers because they exhibited more clear-cut microhabitat preferences: they were almost always hidden into leaf litter of well vegetated, wet and shady spots inside the rainforest, usually in the surroundings of spots with plenty mushrooms. On the other hand, microhabitat characteristics of vipers were less defined (Luiselli 2006a). For the six species of tortoises and snakes, juvenile data were removed from further analyses because of the scarcity of observations at this life stage.

### Goodness-of-fit tests

We examined transience and trap-dependence using U-CARE program (Choquet et al. 2009a). We performed the TEST3.SR, TEST3.SM, TEST2.CT, and TEST2.CL for the six reptile species. The tests TEST3.SR was significant for *K. erosa* and *K. homeana* (two species with a very similar ecology; see Luiselli & Diagne 2013, 2014), which indicates an excess of transients in these species (Supplementary material, Table S5). The other tests were non-significant in all species.

### Modeling survival and recruitment

We examined survival using Cormack-Jolly-Seber models. For tortoises in which an excess of transient was detected we considered a model with three states: transient (T), resident (R), and dead (D). At their first capture, individuals can be in the state T or R. In the following vector probability, individuals may thus be transient with a probability *ψ_T_* or resident with a probability 1 − *ψ_T_*:

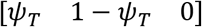

Then, at each time step, resident individuals may survive with a probability *φ_R_* or die with a probability 1 − *φ_R_* while the survival of transient individual is fixed at 0. This results in the following state-state transition matrix (state at time *t*-1 in rows, state at *t* in columns):

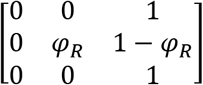

The last component of the model links field observations to underlying states. At each capture session, transient and resident individuals can be captured with a probability *p_T_* and *p_R_*, leading to the following matrix:

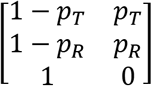

We examined recruitment using Pradel capture-recapture model (1996) in which recruitment is modeled by reversing capture histories and analyzing them backwards. For tortoises, we considered a transient excess and used a modified version of the Pradel model. Recruitment probability was estimated as the probability that an individual present at *t* was not present at *t*-1, i.e. the proportion of “new”, resident individuals in the population at *t*. The model had the same structure than the survival model. However, the survival matrix was replaced by the recruitment matrix. At each time step, resident individuals may be recruited with a probability *δ_T_* or not with a probability 1 − *δ_T_*, leading to the following matrix:

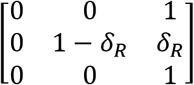

The survival and recruitment models were implemented in program E-SURGE (Choquet et al. 2009b). As the number of years and the study period differed for tortoises and snakes, we analyzed the datasets of tortoises and snakes separately. We ranked models using the second order AIC and its quasi-likelihood counterpart (QAICc) and Akaike weights (w). If the Akaike weight of the best supported model was less than 0.90, we used model-averaging to obtain parameter estimates. The 95% CI were calculated using the delta-method (Royall 1986). For tortoises, we tested our hypotheses about transience, survival, recapture probabilities, from the following general model: [*ψ*(species), *φ*(species), *p*(species + t)] in which species was included as discrete covariate. We also hypothesized that recapture probability varied among years (t). The model for snakes was [*i*(.), *φ*(species), *p*(species + t)]. We tested all the possible combinations of effects, leading to the consideration of 16 and eight competitive models for tortoises and snakes respectively.

For tortoises, we tested our hypotheses about recruitment, transience, and recapture from the model [*ψ*(species), *δ*(species), *p*(species + t)]. For snakes, we considered the model [*i*(.), *δ*(species), *p*(species + t)]. All the possible combinations of effects were considered, leading to 16 and eight candidate models for tortoises and snakes respectively.

### Modeling age-dependent survival and mortality rates

We investigated age-dependent patterns of survival and mortality using Bayesian survival trajectory analyses implemented in the R package BaSTA (Colchero et al. 2012a, 2012b). BaSTA allowed us to account for imperfect detection, left-truncated (i.e., unknown birth date) and right-censored (i.e., unknown death date) capture-recapture data in our analysis. The model allows the estimation of two age-dependent parameters: survival until age *x* and the proportion of individuals dying at age *x* (i.e. mortality rate, or hazard rate).

We focused our analysis at the genus level (the data of *Bitis* species and *Kinixys* species were merged) to increase the statistical power of the analyses. It was not possible to examine age-dependent processes in each species of tortoises and snakes due to the relatively low number of individuals marked; models estimates were too imprecise. We therefore merged the capture-recapture data of the different species of *Kinixys* and *Bitis*. In this regard, it should be considered that *B. gabonica* and *B. nasicornis* are ecologically and morphologically very similar (Luiselli 2006a, 2006b), and the same is true for *K. erosa, K. nogueyi*, and *K. homeana* (Luiselli & Diagne 2013, 2014), thus making our merging of the data as *ecologically* relevant. We analyzed the data of the three genera separately. For tortoises, we removed transient individuals by excluding the first observation in capture-recapture histories. Given the results of the survival models (**Table 1**), we allowed recapture probabilities to vary among years. The proportion of unknown birth date was 0% in *Kinixys* and 13% in *Bitis*, and 12% in *Causus*. The proportion of unknown death date was 4% in *Kinixys*, 0% in *Bitis*, and 1% in *Causus*.

**Table 1.** Model selection procedure for survival models in snakes (*Bitis gabonica, Bitis nasicornis, Causus maculatus*) and tortoises (*Kinixys erosa, Kinixys homeana, Kinixys nogueyi*). r = model rank, k = number of parameters, Dev. = residual deviance, QAICc = quasi-likelihood AICc, ΔQAICc = difference of QAICc points with the best-supported model, w = QAICc weight.

| r | Model | k | Dev. | QAICc | w |
| --- | --- | --- | --- | --- | --- |
| Snake |  |  |  |  |  |
| 1 | $i(\cdot), \varphi(\text{species}), p(\text{species})$ | 6 | 345.56 | 357.94 | 0.59 |
| 2 | $i(\cdot), \varphi(\cdot), p(\text{species})$ | 4 | 351.25 | 359.43 | 0.28 |
| 3 | $i(\cdot), \varphi(\text{species}), p(\cdot)$ | 4 | 353.09 | 361.27 | 0.11 |
| 4 | $i(\cdot), \varphi(\text{species}), p(\text{species}+t)$ | 12 | 340.45 | 365.90 | 0.01 |
| 5 | $i(\cdot), \varphi(\cdot), p(\text{species}+t)$ | 10 | 346.08 | 367.09 | 0.01 |
| 6 | $i(\cdot), \varphi(\text{species}), p(t)$ | 10 | 348.71 | 369.72 | 0.00 |
| 7 | $i(\cdot), \varphi(\cdot), p(\cdot)$ | 2 | 365.37 | 369.42 | 0.00 |
| 8 | $i(\cdot), \varphi(\cdot), p(t)$ | 8 | 360.81 | 377.47 | 0.00 |
| Tortoises |  |  |  |  |  |
| 1 | $\psi(\text{species}), \varphi(\text{species}), p(\text{species}+t)$ | 24 | 1195.63 | 1245.10 | 0.36 |
| 2 | $\psi(\text{species}), \varphi(\cdot), p(\text{species}+t)$ | 22 | 1201.14 | 1246.37 | 0.19 |
| 3 | $\psi(\text{species}), \varphi(\text{species}), p(t)$ | 22 | 1201.93 | 1247.16 | 0.13 |
| 4 | $\psi(\text{species}), \varphi(\text{species}), p(\text{species})$ | 9 | 1228.49 | 1246.71 | 0.16 |
| 5 | $\psi(\text{species}), \varphi(\cdot), p(t)$ | 20 | 1206.99 | 1248.01 | 0.08 |
| 6 | $\psi(\text{species}), \varphi(\cdot), p(\text{species})$ | 7 | 1234.05 | 1248.18 | 0.08 |
| 7 | $\psi(\text{species}), \varphi(\text{species}), p(\cdot)$ | 7 | 1241.77 | 1255.90 | 0.02 |
| 8 | $\psi(\text{species}), \varphi(\cdot), p(\cdot)$ | 5 | 1247.78 | 1257.86 | 0.00 |
| 9 | $\psi(\cdot), \varphi(\cdot), p(t)$ | 18 | 1223.17 | 1260.00 | 0.00 |
| 10 | $\psi(\cdot), \varphi(\cdot), p(\text{species}+t)$ | 20 | 1220.16 | 1261.18 | 0.00 |
| 11 | $\psi(\cdot), \varphi(\text{species}), p(t)$ | 20 | 1221.27 | 1262.30 | 0.00 |
| 12 | $\psi(\cdot), \varphi(\text{species}), p(\text{species}+t)$ | 22 | 1217.88 | 1263.11 | 0.00 |
| 13 | $\psi(\cdot), \varphi(\cdot), p(\text{species})$ | 5 | 1254.33 | 1264.40 | 0.00 |
| 14 | $\psi(\cdot), \varphi(\text{species}), p(\text{species})$ | 7 | 1251.77 | 1265.90 | 0.00 |
| 15 | $\psi(\cdot), \varphi(\text{species}), p(\cdot)$ | 5 | 1262.91 | 1272.98 | 0.00 |
| 16 | $\psi(\cdot), \varphi(\cdot), p(\cdot)$ | 3 | 1267.14 | 1273.16 | 0.00 |

We considered the four mortality functions implemented in BaSTA: exponential, Gompertz, Weibull and logistic. For the three last functions, we considered three potential shapes: *simple* that only uses the basic functions described above; *Makeham* (Pletcher 1999); and *bathtub* (Silver 1979). As individuals usually reach sexual maturity at eight years in *Kinixys* (Coulson & Hailey 2001) and at three years in *Bitis* and *Causus* (Luiselli, unpublished data) genera, we conditioned the analyses at a minimum age of eight in tortoise models and three in snake models. Four MCMC chains were run with 50,000 iterations and a burn-in of 5,000. Chains were thinned by a factor of 50. Model convergence was evaluated using the diagnostic analyses implemented in BaSTA, which calculate the potential scale reduction for each parameter to assess convergence. Models that did not converge were not considered in the procedure of model selection. We used DIC to compare the predictive power of each mortality function and its refinements (Spiegelhalter et al. 2002, Colchero et al. 2012b).

## Results

We made a total of 1071 captures (843 tortoises and 228 snakes). In tortoises, we identified 231 individuals of *K. erosa*, 281 individuals of *K. homeana*, and 57 individuals of *K. nogueyi*. In snakes, we identified 35 individuals of *B. gabonica*, 34 individuals of *B. nasicornis*, and 68 individuals of *C. maculatus*. For more detailed information about capture-recapture data, see m-array matrices in Supplementary material, Table S1-S4.

### Modeling transience, survival and recruitment

For tortoises, the best-supported model was [*ψ*(species), *φ*(species), *p*(species+t)] (**Table 1**); its QAICc weight was 0.36 and we therefore model-averaged the estimates. Recapture probabilities varied according to species and time (Supplementary material, Fig.S1). Recapture probability was the highest in 2013: it was 0.95 (95% 0.82-0.99) in *K. erosa*, 0.90 (95% 0.68-0.98) in *K. homeana*, and 0.90 (95% 0.67-0.97) in *K. nogueyi*. By contrast, recapture probability was the lowest in 2004: it was 0.48 (95% 0.24-0.72) in *K. erosa*, 0.29 (95% 0.14-0.51) in *K. homeana*, and 0.29 (95% 0.10-0.59) in *K. nogueyi*. Moreover, the transience rate differed among species (**Fig.1B**): it was the highest in *K. erosa* (0.79, 95% 0.72-0.85) while *K. homeana* had an intermediate transience rate (0.53, 95% 0.62-0.72), and *K. nogueyi* had the lowest one (0.30, 95% 0.11-0.60). This result is congruent with the GOF tests (Supplementary material, Table S5) that have detected an excess of transience in *K. erosa* and *K. komeana* but not in *K. nogueyi*. Furthermore, survival probability differed between species (**Fig.1A**): *K. erosa* had higher survival (0.83, 95% 0.73-0.90) than *K. homeana* (0.70, 95% 0.62-0.77) and *K. nogueyi* (0.73, 95% 0.56-0.85). The best-supported recruitment model was [*ψ*(species), *δ*(.), *p*(species+t)] (w = 0.23; **Table 2**). Recruitment probability was relatively similar among tortoise species (**Fig.1C**): it was 0.28 (95% 0.22-0.35) in *K. erosa*, 0.28 (95% 0.22-0.34) in *K. homeana*, and 0.33 (95% 0.25-0.42) in *K. nogueyi*.

**Fig.1.**
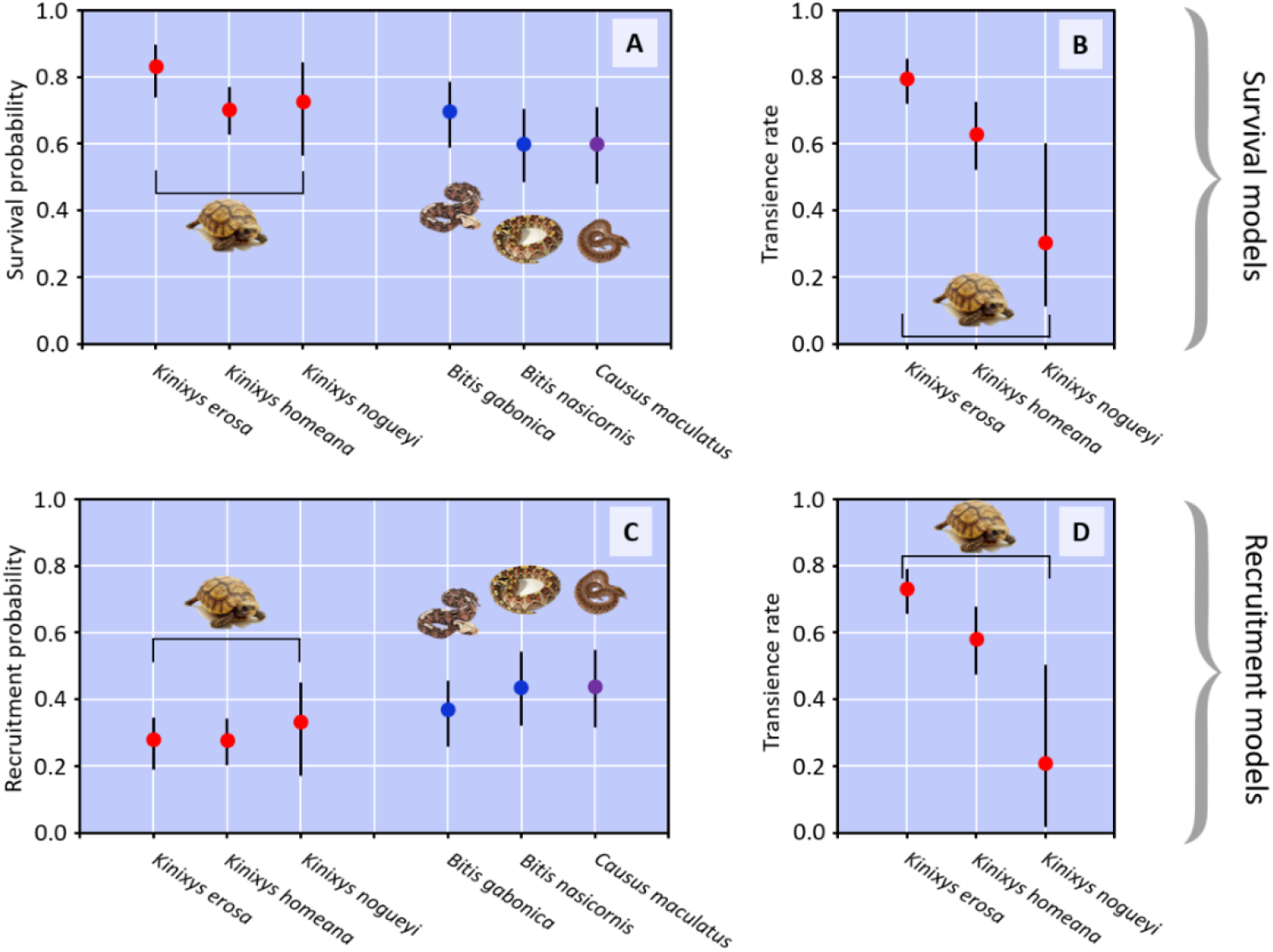
Estimates of survival, recruitment, and transience in three tropical tortoises (*Kinixys erosa, Kinixys homeana, Kinixys nogueyi*) and snakes (*Bitis gabonica, Bitis nasicornis, Causus maculatus*).

**Table 2.** Model selection procedure for recruitment models in snakes (*Bitis gabonica, Bitis nasicornis, Causus maculatus*) and tortoises (*Kinixys erosa, Kinixys homeana, Kinixys nogueyi*). r = model rank, k = number of parameters, Dev. = residual deviance, QAICc = quasi-likelihood AICc, ΔQAICc = difference of QAICc points with the best-supported model, w = QAICc weight.

| r | Model | k | Dev. | QAICc | w |
| --- | --- | --- | --- | --- | --- |
| Snakes |  |  |  |  |  |
| 1 | $i(\cdot), \delta(\cdot), p(\text{species})$ | 4 | 371.62 | 379.80 | 0.55 |
| 2 | $i(\cdot), \delta(\text{species}), p(\text{species})$ | 6 | 368.47 | 380.85 | 0.32 |
| 3 | $i(\cdot), \delta(\text{species}), p(\cdot)$ | 4 | 375.10 | 383.28 | 0.10 |
| 4 | $i(\cdot), \delta(\cdot), p(\cdot)$ | 2 | 382.49 | 386.55 | 0.02 |
| 5 | $i(\cdot), \delta(\cdot), p(\text{species}+t)$ | 10 | 368.29 | 389.31 | 0.00 |
| 6 | $i(\cdot), \delta(\text{species}), p(\text{species}+t)$ | 12 | 365.25 | 390.70 | 0.00 |
| 7 | $i(\cdot), \delta(\text{species}), p(t)$ | 10 | 371.60 | 392.62 | 0.00 |
| 8 | $i(\cdot), \delta(\cdot), p(t)$ | 8 | 378.87 | 395.53 | 0.00 |
| Tortoises |  |  |  |  |  |
| 1 | $\psi(\text{species}), \delta(\cdot), p(\text{species}+t)$ | 22 | 1254.18 | 1299.41 | 0.23 |
| 2 | $\psi(\text{species}), \delta(\cdot), p(\text{species})$ | 7 | 1284.84 | 1298.98 | 0.28 |
| 3 | $\psi(\text{species}), \delta(\text{species}), p(\text{species}+t)$ | 24 | 1251.52 | 1300.98 | 0.10 |
| 4 | $\psi(\text{species}), \delta(\text{species}), p(\text{species})$ | 9 | 1281.62 | 1299.83 | 0.18 |
| 5 | $\psi(\text{species}), \delta(\cdot), p(t)$ | 20 | 1259.64 | 1300.66 | 0.12 |
| 6 | $\psi(\text{species}), \delta(\text{species}), p(t)$ | 22 | 1256.39 | 1301.62 | 0.07 |
| 7 | $\psi(\text{species}), \delta(\cdot), p(\cdot)$ | 5 | 1296.18 | 1306.25 | 0.01 |
| 8 | $\psi(\text{species}), \delta(\text{species}), p(\cdot)$ | 7 | 1293.03 | 1307.17 | 0.00 |
| 9 | $\psi(\cdot), \delta(\cdot), p(t)$ | 18 | 1275.02 | 1311.85 | 0.00 |
| 10 | $\psi(\cdot), \delta(\text{species}), p(t)$ | 20 | 1273.69 | 1314.71 | 0.00 |
| 11 | $\psi(\cdot), \delta(\cdot), p(\text{species}+t)$ | 20 | 1273.83 | 1314.85 | 0.00 |
| 12 | $\psi(\cdot), \delta(\cdot), p(\text{species})$ | 5 | 1304.33 | 1314.40 | 0.00 |
| 13 | $\psi(\cdot), \delta(\text{species}), p(\text{species}+t)$ | 22 | 1272.06 | 1317.30 | 0.00 |
| 14 | $\psi(\cdot), \delta(\text{species}), p(\text{species})$ | 7 | 1302.66 | 1316.79 | 0.00 |
| 15 | $\psi(\cdot), \delta(\cdot), p(\cdot)$ | 3 | 1314.02 | 1320.05 | 0.00 |
| 16 | $\psi(\cdot), \delta(\text{species}), p(\cdot)$ | 5 | 1313.33 | 1323.40 | 0.00 |

For snakes, [*i*(.), *φ*(species), *p*(species)] was the best supported model (w = 0.59, **Table 1**). The recapture probability marginally varied among years (Supplementary material, Fig.S1). However, it markedly differed among species: *C. maculatus* had a higher recapture probability (e.g., 2003: 0.93, 95% CI 0.67-0.99) than *B. gabonica* (2003: 0.83, 95% CI 0.56-0.95) and *B. nasicornis* (2003: 0.61, 95% CI 0.29-0.85). Moreover, survival rate slightly varied among species: *B. gabonica* (0.70, 95% CI 0.59-0.78) had a higher survival rate than *B. nasicornis* (0.60, 95% CI 0.48-0.70) and *C. maculatus* (0.61, 95% CI 0.49-0.71). The best-supported recruitment model was [*i*(.), *δ*(.), *p*(species+t)] (w = 0.23; **Table 2**). Recruitment probability was relatively similar among snake species (**Fig.1C**): it was 0.37 (95% 0.29-0.47) in *B. gabonica*, 0.44 (95% 0.33-0.55) in *B. nasicornis*, and 0.44 (95% 0.33-0.55) in *C. maculatus*.

### Modeling age-dependent survival and mortality rates

In the genus *Kinixys*, data were best described by a Weibull function (Supplementary material, Table S2). The model indicated the mortality rate was age-dependent: it increased gradually with age (**Fig.2A**). Survival probability was 0.95 until nine years, 0.50 until 12 years, 0.25 until 13 years, and was below than 0.05 until 16 years (**Fig.2D**).

**Fig.2.**
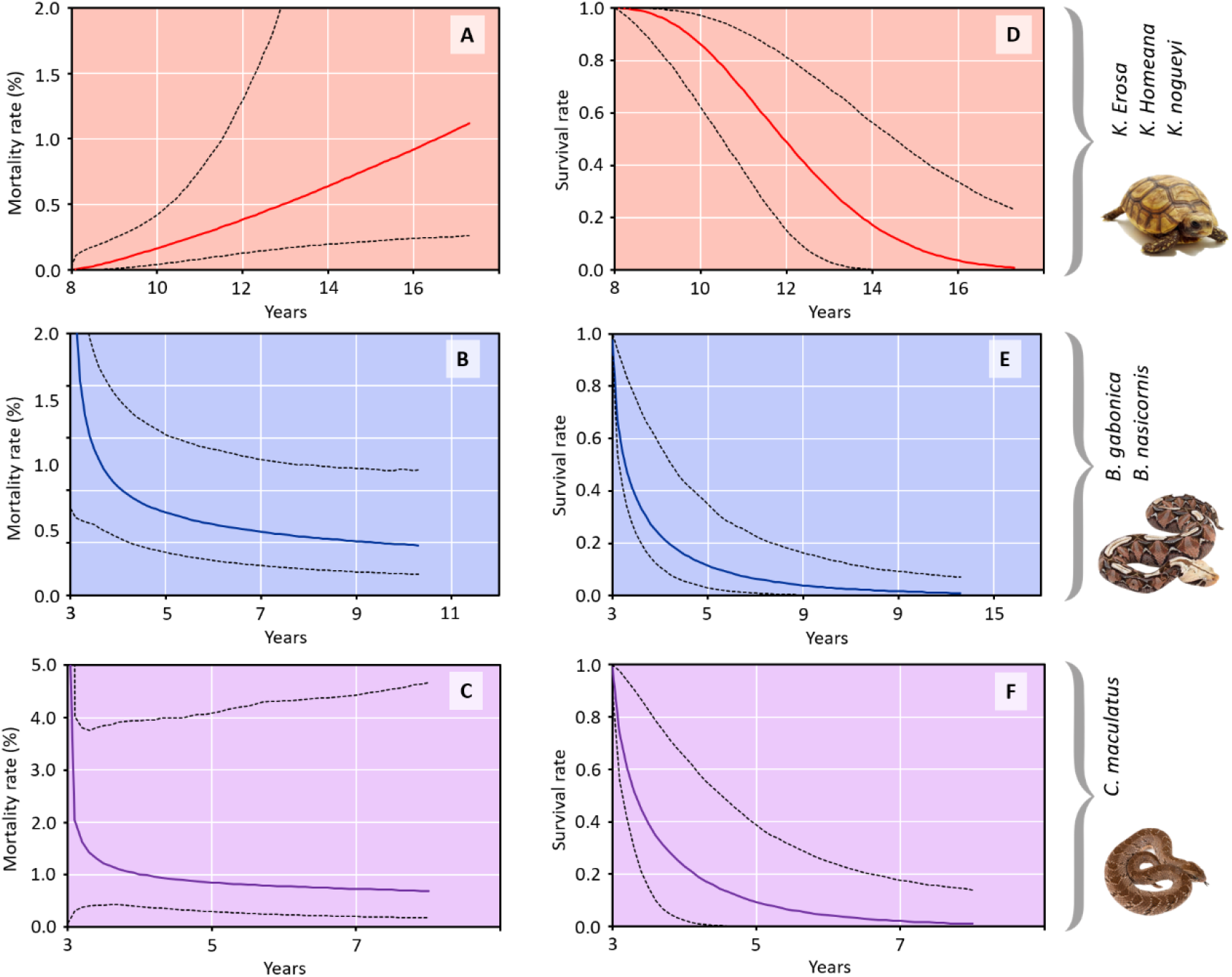
Age-dependent survival and mortality (i.e., hazard) rate in the genera *Kinixys, Bitis*, and *Causus*.

As well, the Weibull function was the best supported one in the genus *Bitis* (Supplementary material, Table S2). The model showed that mortality rates decreased with age (**Fig.2B**): it dropped dramatically and then the decrease tended to be less strong. Survival probability was 0.95 until 3 years, 0.50 until 3.5 years, 0.25 until 4 years, and was below than 0.05 until 7 years (**Fig.2E**).

In the genus *Causus*, exactly as in the two previous genera, data were best described by a Weibull function (Supplementary material, Table S2). Mortality rate was age-dependent although the 95% CI were very large (**Fig.2B**). Mortality rate decreased brutally and then remained stable. Survival probability was 0.95 until three years, 0.50 at 3.5 years, 0.25 until 4 years, and was below than 0.05 until 5.5 years (**Fig.2E**).

## Discussion

We validated two out of our three hypotheses of departure. First, we showed that tortoises of the *Kinixys* genus had a higher survival and a lower recruitment than snakes of the genera *Bitis* and *Causus*, indicating that they have a slower life history (*hypothesis 1*). Second, we showed that survival more slowly decreased with age in tortoises than in snakes (*hypothesis 2*). Third, we highlighted contrasted age-dependent mortality rate patterns in the three genera. In *Kinixys*, the relationship between mortality rate and age was positive and linear, suggesting gradual senescence over tortoise lifetime. By contrast, the relationship between mortality and age was negative and sharp in *Bitis* and *Causus*, suggesting negative senescence starting early in life. Therefore, we did not validated *hypothesis 3* (i.e., a more delayed senescence in *Kinixys* with slow life histories) as only tortoises likely experienced “positive” senescence.

### Population dynamics and species position along the fast-slow continuum

The tortoises of the genus *Kinixys* had slower life histories (i.e. longer lifespan and lower recruitment) than the three snake species. The total lifespan obtained by adding the pre-maturity lifespan (8 years, Lawson 2001) and the adult lifespan (calculated using survival *φ* estimates; lifespan = 1-/ln(*φ*)+ 8) was 13 years in *K. erosa*, and 11 years in *K. homeana* and *K. nogueyi*. Those estimates are congruent with the lifespan (10 years) calculated using scale ring counting in *Kinixys spekii* (Coulson & Hailey 2001). In snakes, the total lifespan was 6 years in *B. gabonica* and 5 years in *B. nasicornis* and *C. maculatus*. The lifespan of the *Bitis* species is far lower than the one reported in captivity (around 18 years). First, this could be due to methodological limitations: in capture-recapture studies, survival can be biased by permanent emigration from the study area (Lebreton et al. 1992). Large *Bitis* species such as *B. gabonica* or *B. nasicornis* can exhibit looping excursions well outside their home ranges (Linn et al. 2006), which may lead to apparent survival if they die before returning to their home range. Yet, this explanation does not seem completely satisfactory as recapture probability is relatively high in *B. gabonica* and GOF test 3SR did not indicate an excess of transient, which does not suggest a high permanent emigration from the study area. Alternatively, the low survival of snakes may be due to a high mortality in natural conditions. Small adults could experience predation but anthropogenic factors (road killing, voluntary destruction due to snake harmfulness, human hunting for subsistence) might also negatively affect survival. Indeed, *Bitis gabonica* is one of the most intensely hunted snakes in the Niger Delta region for the bushmeat trade (Eniang et al. 2006, Akani et al. unpublished data). Since these vipers are actively searched for by hunters, it is likely that their mortality risks are high (given also the high density of settlements and population around the forested patches), and this might have substantially reduced the life expectancy of these vipers locally. For *B. nasicornis* it is the same, but this latter species occurs less frequently than *B. gabonica* in the local bushmeat markets (Eniang et al. 2006, Akani et al. unpublished data).

In parallel, recruitment was lower in tortoises than in snakes. Recruitment probability was relatively similar (around 0.30) among the tortoises of the *Kinixys* genus. By contrast, the recruitment probability was slightly higher in the three snake species (around 0.40) and did not markedly differed between the genera *Bitis* and *Causus*. The lower recruitment in *Kinixys* than in *Bitis* and *Causus* likely resulted from a variation in female fecundity. Females of *K. erosa* and *K. homeana* lay from 4 and 8 eggs (Akani et al. 2004) while females of *B. gabonica* and *B. nasicornis* produce 18 and 25 young respectively after a gestation of one year.

### Footprints of positive and negative senescence

We only detected the footprint of (positive) senescence in tortoises. In the *Kinixys* genus, individuals appear to experience a gradual senescence: the relationship between mortality rate and age was almost linear and reached 100% at 17 years. This pattern markedly differs from the one reported (i.e. negative senescence) by Jones et al. (2014) for the desert tortoise *Gopherus agassizii*. However, *Kinixys* and *Gopherus* have very contrasted ecological characteristics: *Gopherus* are burrowers, vegetarian species from dry, moderately vegetated up to semidesertic areas (e.g. Ashton & Ashton, 2007), whereas *Kinixys* are above-ground active, omnivorous species from very wet, forested areas (Luiselli & Diagne 2013, 2014). In addition, age-dependent survival patterns strongly differ between *Kinixys* and *Gopherus* tortoises. In *Gopherus agassizii*, juveniles experience a high mortality while adult have very high survival (*φ*> 0.95, Tuberville et al. 2008), which results in a very long lifespan (around 40 years, Curtin et al. 2009) in few individuals and a negative senescence pattern. By contrast, our study and a previous one (Coulson & Hailey 2001) indicated that *Kinixys* tortoises have a shorter lifespan (13-10 years) likely associated with a progressive senescence over individual lifetime.

The age-dependent mortality pattern found in *Bitis* and *Causus* vipers strongly suggests a negative senescence (Vaupel et al. 2004). In the genus *Bitis*, the mortality rate sharply decreases between three and five years and then tends to slow down. In *Causus*, mortality dramatically dropped between three and four years and then stabilized. This indicates that vipers experience a high mortality during a period (few years) following sexual maturity. Mortality tends to decrease after that allowing few individuals to have a relatively long lifespan (possibly more than 10 years). In those snakes, negative senescence likely results from large variation of body size over snake lifetime and size-dependent survival. In *B. gabonica* for instance, newborns have a body size of 0.30 m and body mass of 0.05 kg while large adults can reach a size of 2.2 meters and a mass of 10 kg (Bonnet et al. 2001). It is possible that young adults (3-4 years) with a relatively small body size (0.80-1.10 m) experience a high mortality due to extrinsic factors such as predation while a large size may better protect old individuals from those factors. Cobras frequently eat on vipers and other snakes (Luiselli et al. 2002, Filippi & Petretto 2013, Maritz et al. 2019) and two species, *Naja melanoleuca* and *Naja nigricollis*, are common at the study area. The pattern of negative senescence found in *Bitis* and *Causus* strongly differs from the one reported for *Vipera aspis*, a small viperid from temperate regions that seems to do not experience either positive or negative senescence (Colchero et al. 2019).

## Conclusion

To our knowledge, the present study was the first to investigate age-dependent mortality processes in tropical squamates and contributed to extend our knowledge of senescence in amniotes in a more general way. Our results, and those of Jones et al. (2014), indicate that negative senescence that was initially ruled out by the Hamilton’s model (Hamilton 1966) seems to be a common pattern in reptiles while it has not been reported so far in mammals and birds. They also indicate that reptiles with contrasted life histories and population dynamics may have highly divergent senescence patterns. We strongly encourage further studies to use capture-recapture data available in a broader range of ectotherm amniotes to expand our understanding of senescence in the living world.

## Acknowledgements

Funding of *Kinixys* research (that also supported the snake study) over the years were provided by two grants from the Mohamed Bin Zayed Species Conservation Fund, five grants from the Turtle Conservation Fund, three “Linnaeus Funds” grants from the Chelonian Research Foundation, and by one grant by the Andrew Sabin and Family Foundation (all funds to LL as lead researcher). We are indebted with the Federal Department of Forestry (Port Harcourt) for authorization to perform the study, and to the Rivers State University of Science and Technology (Port Harcourt) and the University of Uyo for logistic assistance.

## Supplementary material

**Table S1.**
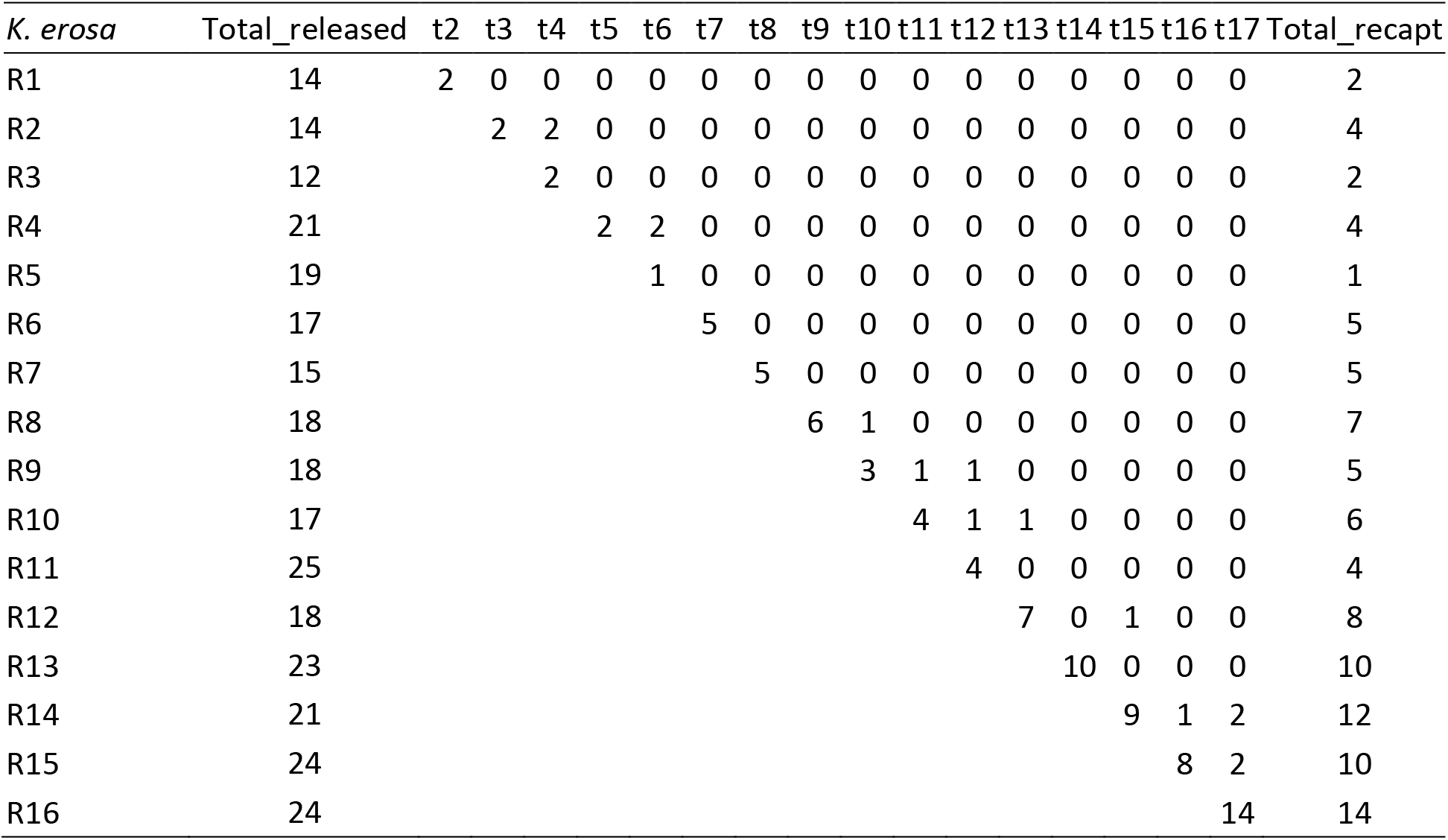
M-array matrix for *K. erosa*.

**Table S2.**
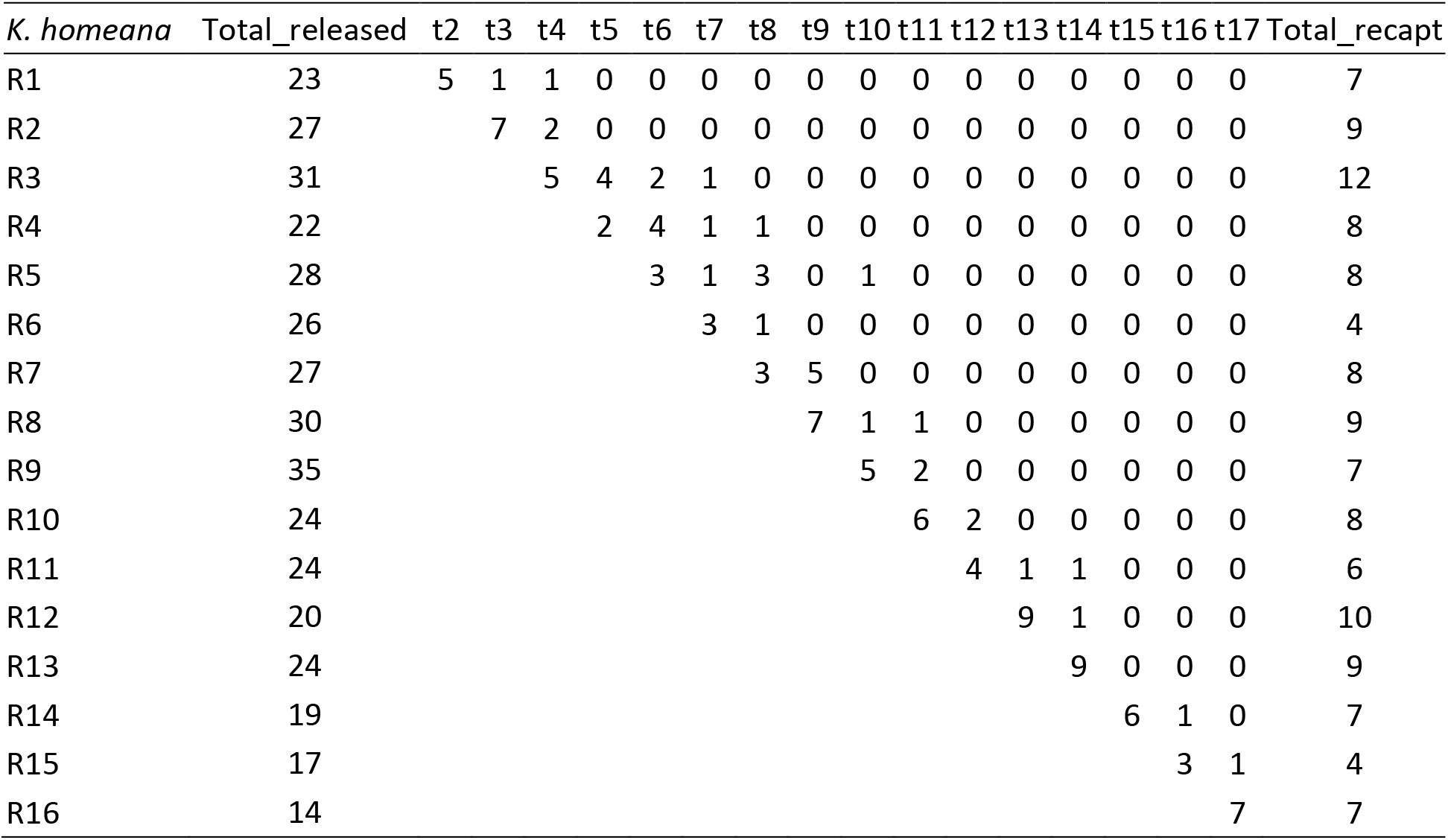
M-array matrix for *K. homeana*.

**Table S3.**
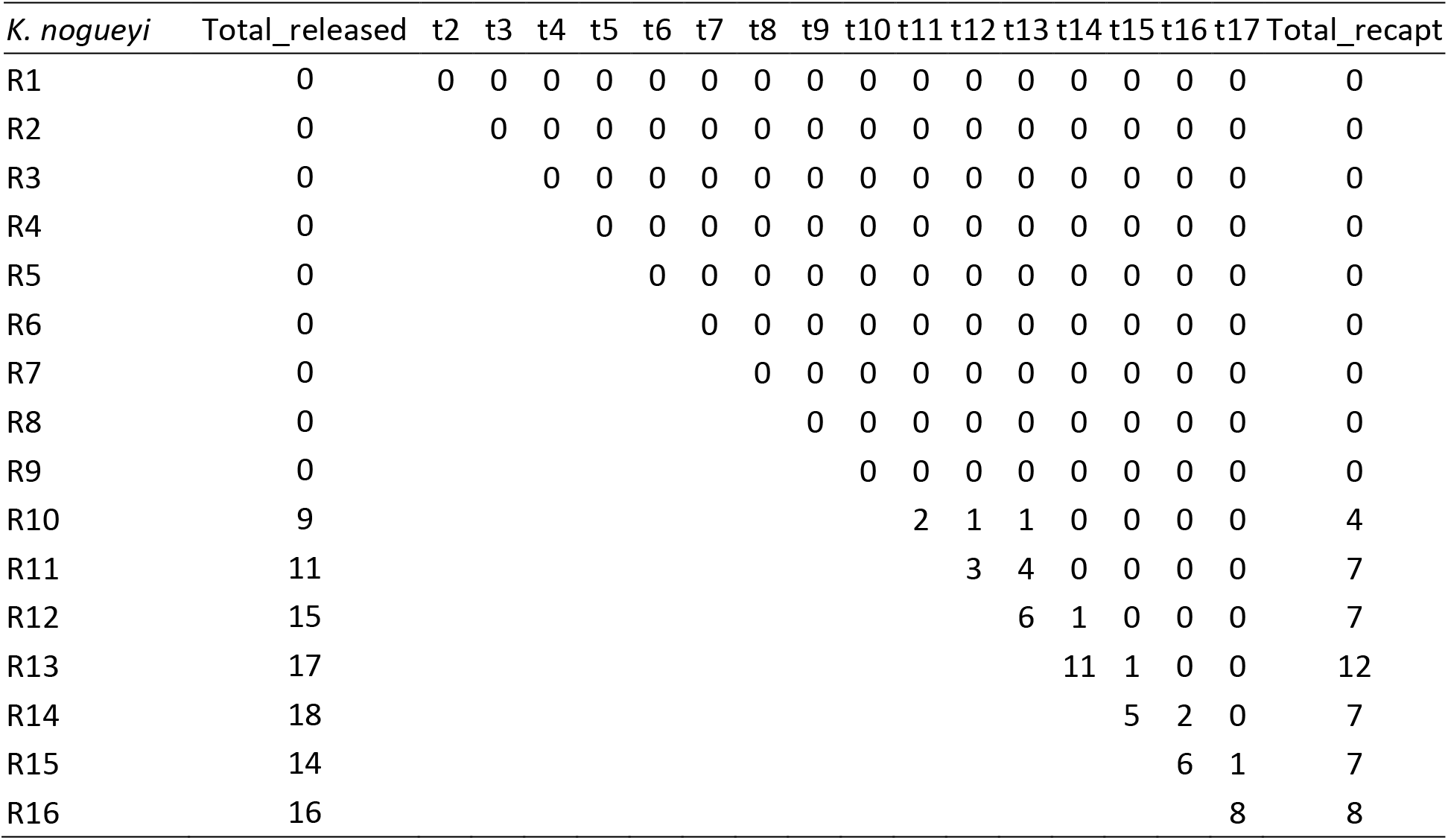
M-array matrix for *K. nogueyi*.

**Table S4.**
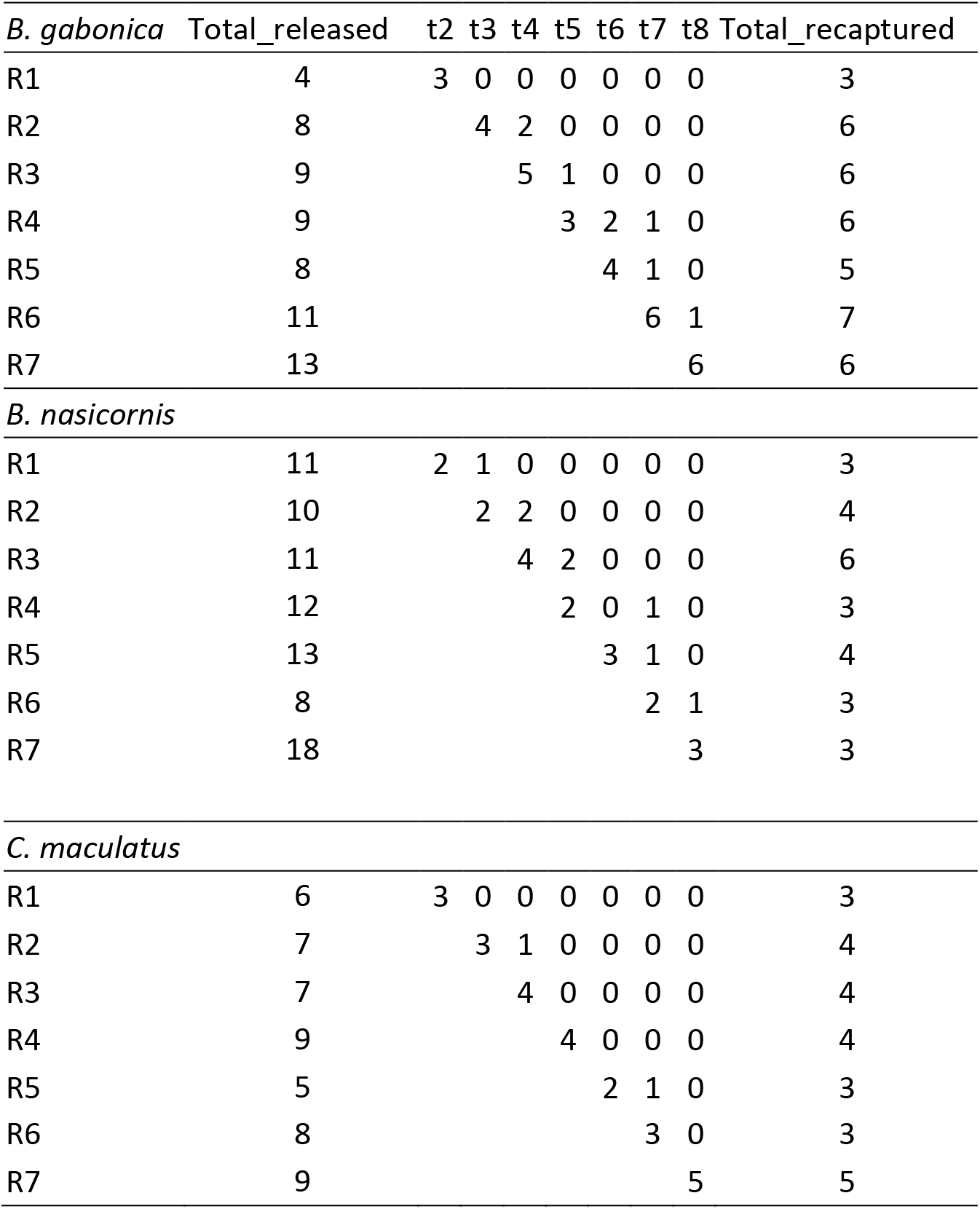
M-array matrix for the three snake species.

**Table S5.**
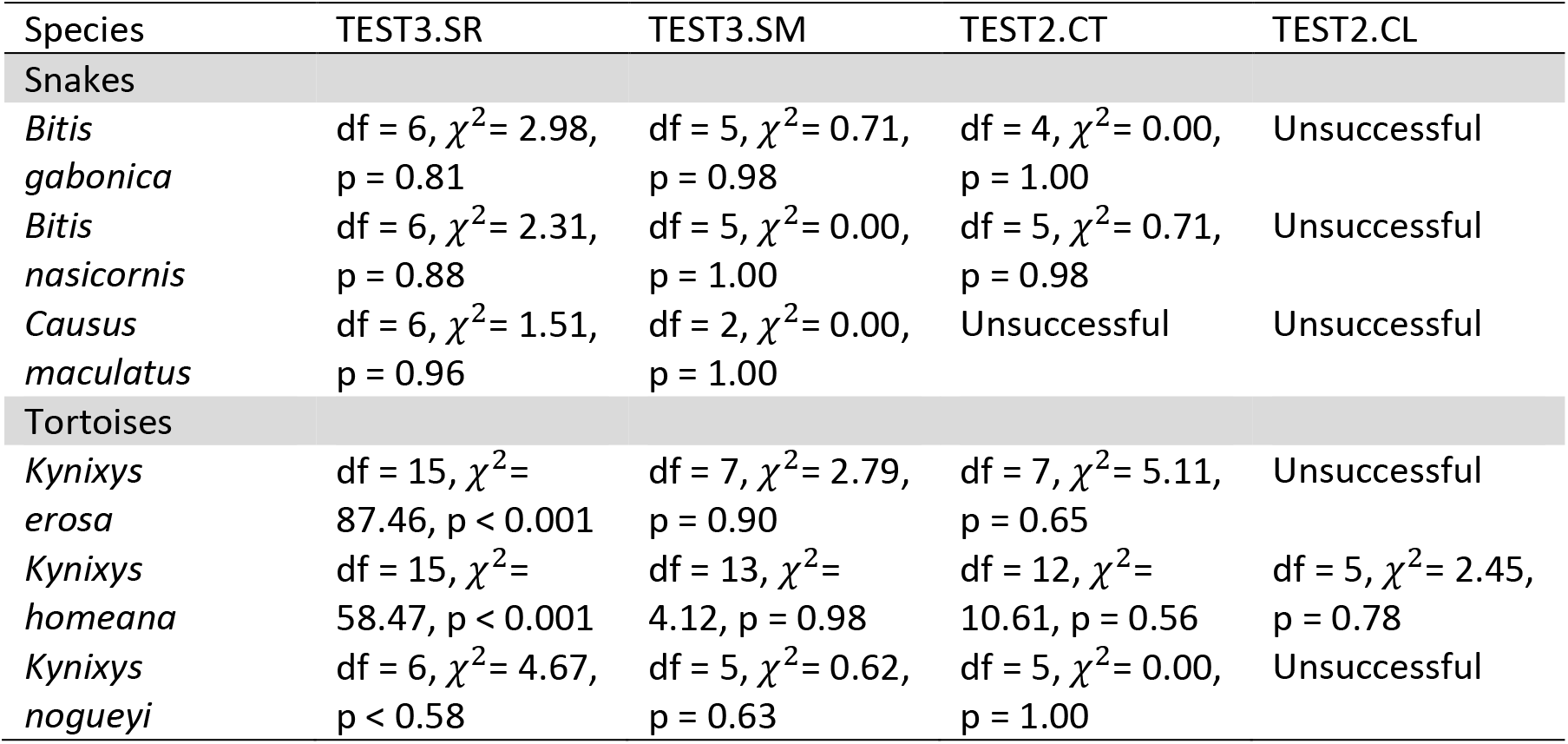
Goodness-of-fit tests (TEST3.SR, TEST3.SM, TEST2.CT, and TEST2.CL) performed in the program U-CARE for the six species of reptiles.

**Table S6.** Age-dependent survival and mortality rate in the genus *Kinixys*. Deviance information criterion (DIC) for each of the mortality function. We considered the four mortality functions implemented in BaSTA program: exponential (EXP), Gompertz (GOM), Weibull (WEI) and logistic (LOG). For the three last functions, we considered three potential shapes: simple that only uses the basic functions described above (“simple”); Makeham (“make”); and bathtub (“bath”).

|  | *Simple* | *Bath* | *Makeham* |
| --- | --- | --- | --- |
| Logistic | 1660.15 | 1648.95 | 1657.82 |
| Exponential | 1753.58 | - | - |
| Gompertz | Not converged | Not converged | 1640.35 |
| Weibull | 1631.98 | 1652.27 | 1668.54 |

**Table S7.** Age-dependent survival and mortality rate in the genus *Bitis*.

|  | *Simple* | *Bath* | *Makeham* |
| --- | --- | --- | --- |
| Logistic | 601.03 | Not converged | 611.46 |
| Exponential | 599.11 | - | - |
| Gompertz | Not converged | Not converged | Not converged |
| Weibull | 584.10 | 592.71 | Not converged |

**Table S8.** Age-dependent survival and mortality rate in the genus *Causus*.

|  | *Simple* | *Bath* | *Makeham* |
| --- | --- | --- | --- |
| Logistic | 402.59 | Not converged | Not converged |
| Exponential | 370.63 | - | - |
| Gompertz | Not converged | Not converged | Not converged |
| Weibull | 231.96 | 238.54 | Not converged |

**Fig.S1.**
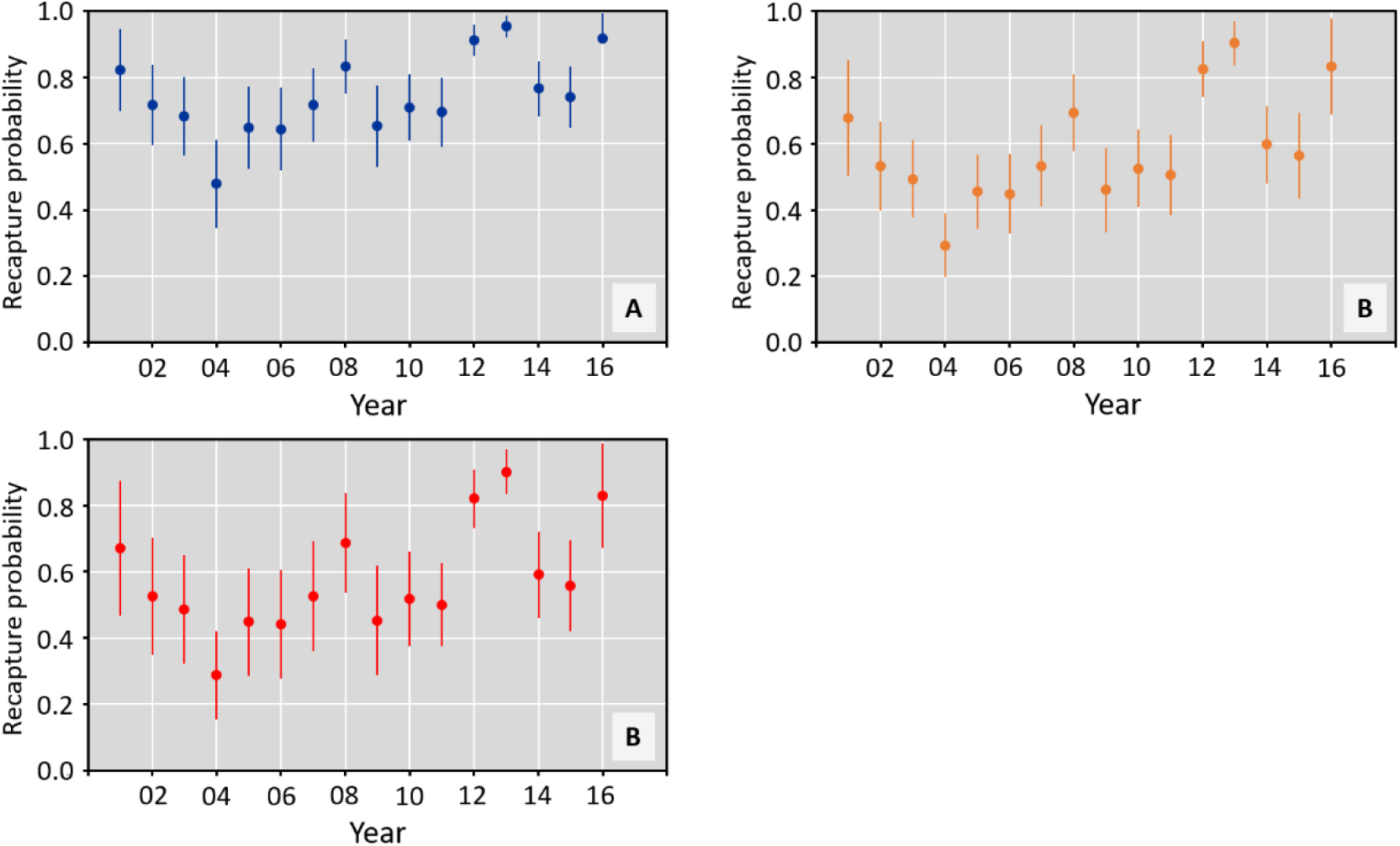
Recapture probability of *Kinixys erosa, Kinixys homeana*, and *Kinixys nogueyi* over the period 2001-2016. Model-averaged estimates and standard errors.

**Fig.S2.**
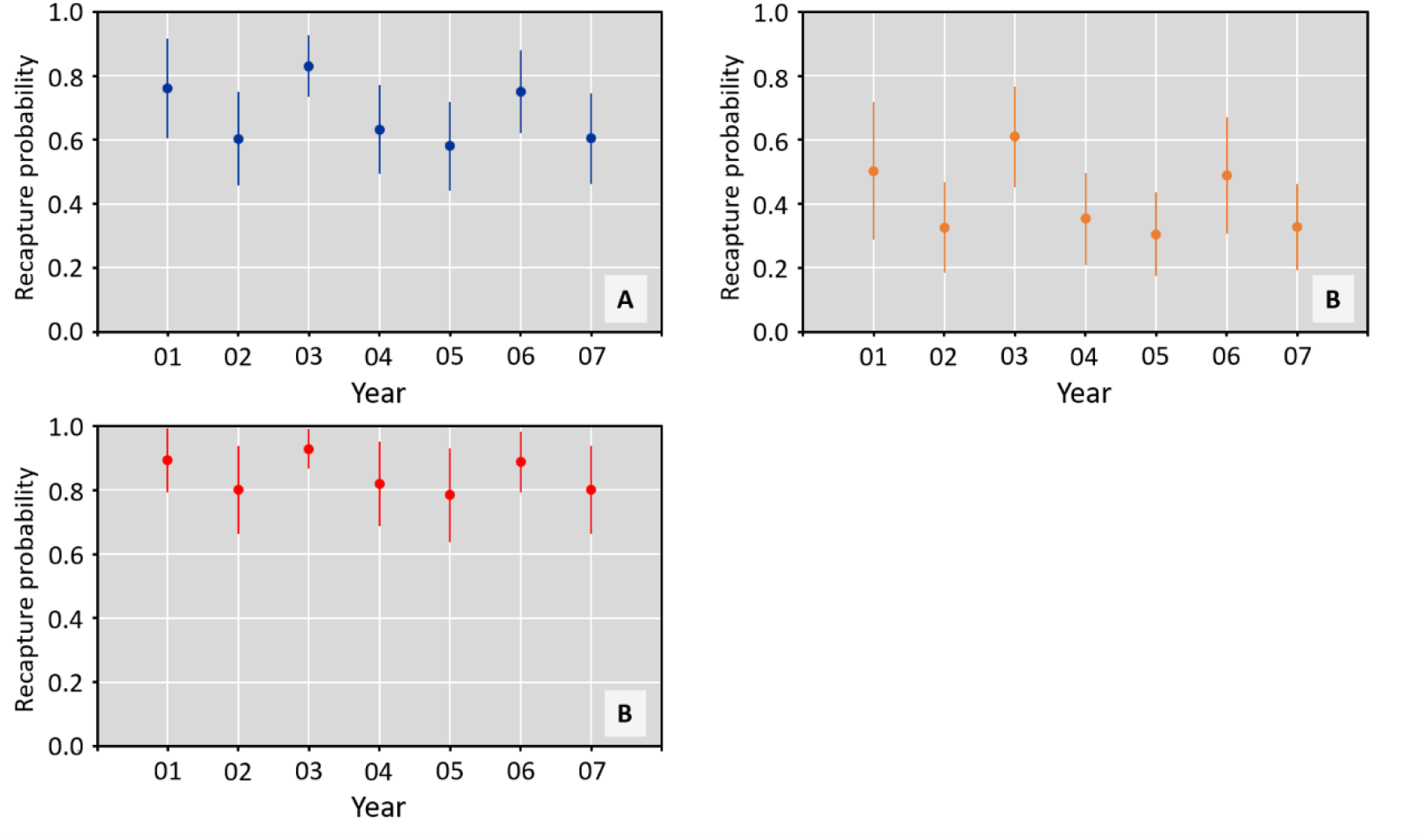
Recapture probability of *Bitis gabonica, Bitis nasicornis*, and *Causus maculatus* over the period 2001-2007. Model-averaged estimates and standard errors.

## References

Akani, G.C., Dendi, D., & Luiselli, L. (2015) Ebola virus effects on the bushmeat trade in West Africa. African Journal of Ecology, 53, 613–615.

Akani, G. C., Filippi, E., & Luiselli, L. (2004). Aspects of the population and reproductive ecology of sympatric hinge-back tortoises (*Kinixys homeana* and *K. erosa*) in southern Nigeria, on the basis of specimens traded in bush-meat markets. Italian Journal of Zoology, 71, 245–247.

Ashton, R.E., & Ashton, P.S. (2007) The Natural History and Management of the Gopher Tortoise Gopherus polyphemus (Daudin). Krieger Publishing Company, Malabar, US.

Baudisch, A., Salguero-Gómez, R., Jones, O. R., Wrycza, T., Mbeau-Ache, C., Franco, M., & Colchero, F. (2013). The pace and shape of senescence in angiosperms. Journal of Ecology, 101, 596–606.

Bielby, J., Mace, G. M., Bininda-Emonds, O. R., Cardillo, M., Gittleman, J. L., Jones, K. E., Orme, C. D. L, & Purvis, A. (2007). The fast-slow continuum in mammalian life history: an empirical reevaluation. The American Naturalist, 169, 748–757.

Bonnet, X., Shine, R., Naulleau, G., & Thiburce, C. (2001). Plastic vipers: influence of food intake on the size and shape of Gaboon vipers (Bitis gabonica). Journal of Zoology, 255, 341–351.

Chippaux, J.-P. (2013). Les serpents d’Afrique occidentale et centrale. IRD Editions, Paris, France.

Choquet, R., Lebreton, J. D., Gimenez, O., Reboulet, A. M., & Pradel, R. (2009a). U-CARE: Utilities for performing goodness of fit tests and manipulating CApture–REcapture data. Ecography, 32, 1071–1074.

Choquet, R., Rouan, L., & Pradel, R. (2009). Program E-SURGE: a software application for fitting multievent models. In: Thomson, D. L., Cooch, E. G., Conroy, M. J. (eds). Modeling demographic processes in marked populations. Pp. 845–865. Springer, Boston, US.

Colchero, F., Jones, O. R., Conde, D. A., Hodgson, D., Zajitschek, F., Schmidt, B. R., Malo, A. F., Alberts, S. C., Becker, P. H., Bouwhuis, S. Bronikowski, A. M., De Vleeschouwer, K. M., Delahay, R. J., Dummermuth, S., Fernández-Duque, E., Frisenvænge, J., Hesselsøe, M., Larson, S., J.-F. Lemaître, McDonald, J., Miller, D. A. W., O’Donnell, C., Packer, C., Raboy, B. E., Reading, C. J., Wapstra, E., Weimerskirch, H., While, G. M., Baudisch, A., Flatt, T., Coulson, T., & Gaillard, J.-M. (2019). The diversity of population responses to environmental change. Ecology Letters, 22, 342–353.

Congdon, J. D., Dunham, A. E., & van Loben Sels, R. C. (1993). Delayed sexual maturity and demographics of Blanding’s turtles (*Emydoidea blandingii*): implications for conservation and management of long-lived organisms. Conservation Biology, 7, 826–833.

Congdon, J. D., Dunham, A. E., & Sels, R. V. L. (1994). Demographics of common snapping turtles (*Chelydra serpentina*): implications for conservation and management of long-lived organisms. American Zoologist, 34, 397–408.

Coulson, I. M., & Hailey, A. (2001). Low survival rate and high predation in the African hingeback tortoise *Kinixys spekii*. African Journal of Ecology, 39, 383–392.

Curtin, A. J., Zug, G. R., & Spotila, J. R. (2009). Longevity and growth strategies of the desert tortoise (*Gopherus agassizii*) in two American deserts. Journal of Arid Environments, 73, 463–471.

Eniang, E.A., Egwali, E.C., Luiselli, L.M., Ayodele, I.A., Akani, G.C., Pacini, N. (2006) Snake bushmeat from the forest markets of south-eastern Nigeria. Natura, 95, 33–46.

Filippi, E., Petretto, M. (2013) *Naja haje* (Egyptian cobra) Diet / ophiophagy. Herpetological Review, 44, 155–156.

Fridlyanskaya, I., Alekseenko, L., & Nikolsky, N. (2015). Senescence as a general cellular response to stress: a mini-review. Experimental Gerontology, 72, 124–128.

Gibbons, J. W. (1987). Why do turtles live so long?. BioScience, 37, 262–269.

Hamilton, W.D. (1966) Moulding of senescence by natural selection. Journal of Theoretical Biology, 12, 12–45.

Jones O. R., Gaillard J.-M., Tuljapurkar S., Alho J. S., Armitage K. B., Becker P. H., Bize P., Brommer, J., Charmantier, A., Charpentier, M., Clutton-Brock, T., Dobson, F. S., Festa-Bianchet, M., Gustafsson, L., Jensen, H, Jones, C. G., Lillandt, B. G., Mc Cleery, R., Merilä, J., Neuhaus, P., Nicoll, M. A. C., Norris, K., Oli, M. K., Pemberton, J., Pietiäinen, H., Ringsby, T. H., Roulin, A., Saether, B. E., Setchell, J. M., Sheldon, B. C., Thompson, P. M., Weimerskirch, H., Wickings, E. J., & Coulson, T. (2008). Senescence rates are determined by ranking on the fast–slow life-history continuum. Ecology Letters, 11, 664–673.

Jones, O. R., Scheuerlein, A., Salguero-Gómez, R., Camarda, C. G., Schaible, R., Casper, B. B., Dahlgren, J. P., Ehrlén, J., García, M. B., Menges, E. S., Quintana-Ascencio, P. F., Caswell, H., Baudisch, A., & Vaupel, J. W. (2014). Diversity of ageing across the tree of life. Nature 505, 169–173.

Jones, O. R., & Vaupel, J. W. (2017). Senescence is not inevitable. Biogerontology, 18, 965–971.

Kiørboe, T., Ceballos, S., & Thygesen, U. H. (2015). Interrelations between senescence, life-history traits, and behavior in planktonic copepods. Ecology, 96, 2225–2235.

Kirkwood, T.B.L. (1977) Evolution of aging. Nature, 270, 301–304.

Lawson, D. P. (2001). Morphometrics and sexual dimorphism of the hinge-back tortoises *Kinixys erosa* and *Kinixys homeana* (Reptilia: Testudinidae) in southwestern Cameroon. African Journal of Herpetology, 50, 1–7.

Lebreton, J. D., Burnham, K. P., Clobert, J., & Anderson, D. R. (1992). Modeling survival and testing biological hypotheses using marked animals: a unified approach with case studies. Ecological Monographs, 62, 67–118.

Linn, I. J., Perrin, M. R., Bodbijl, T. (2006) Movements and home range of the Gaboon adder, *Bitis gabonica gabonica*, in Zululand, South Africa. African Zoology, 41, 252–265.

Luiselli, L. (2003a) Seasonal activity patterns and diet divergence of three sympatric Afrotropical tortoise species (genus *Kinixys*). Contributions to Zoology, 72, 211–220.

Luiselli, L. (2003b) Assessing the impact of human hunting activities on populations of forest tortoises (genus *Kinixys*) in the Niger Delta, Nigeria. Chelonian Conservation and Biology, 4, 735–738.

Luiselli, L. (2006a) Site occupancy and density of sympatric Gaboon viper (*Bitis gabonica*) and nose-horned viper (*Bitis nasicornis*). Journal of Tropical Ecology, 22, 555–564.

Luiselli, L. (2006b) Food niche overlap between sympatric potential competitors increases with habitat alteration at different trophic levels in rain-forest reptiles (omnivorous tortoises and carnivorous vipers). Journal of Tropical Ecology, 22, 695–704.

Luiselli, L., Angelici, F.M., Akani, G.C. (2002) Comparative feeding strategies and dietary plasticity of the sympatric cobras *Naja melanoleuca* and *Naja nigricollis* in three diverging Afrotropical habitats. Canadian Journal of Zoology, 80, 55–63.

Luiselli, L., Akani, G. C. (2003). Diet of sympatric Gaboon vipers (*Bitis gabonica*) and nose-horned vipers (Bitis nasicornis) in southern Nigeria. African Journal of Herpetology, 52, 101–106.

Luiselli, L. and Diagne, T. (2013) *Kinixys homeana* Bell 1827 – Home’s Hinge-Back Tortoise. In: Rhodin, A.G.J., Pritchard, P.C.H., van Dijk, P.P., Saumure, R.A., Buhlmann, K.A., Iverson, J.B., and Mittermeier, R.A. (Eds.). Conservation Biology of Freshwater Turtles and Tortoises: A Compilation Project of the IUCN/SSC Tortoise and Freshwater Turtle Specialist Group. Chelonian Research Monographs, 5, 070.1–070.10.

Luiselli, L. and Diagne, T. (2014) *Kinixys erosa* (Schweigger 1812) – Forest Hinge-back Tortoise, Serrated Hinge-back Tortoise, Serrated Hinged Tortoise. In: Rhodin, A.G.J., Pritchard, P.C.H., van Dijk, P.P., Saumure, R.A., Buhlmann, K.A., Iverson, J.B., and Mittermeier, R.A. (Eds.). Conservation Biology of Freshwater Turtles and Tortoises: A Compilation Project of the IUCN/SSC Tortoise and Freshwater Turtle Specialist Group. Chelonian Research Monographs, 5, 084.1–13.

Luiselli, L., Hema, E.M., Segniagbeto, G.H., Ouattara, V., Eniang, E.A., Parfait, G., Akani, G.C., Sirima, D., Fakae, B.B., Dendi, D., Fa, J.E. (2018) Bushmeat consumption in large urban centres in West Africa. Oryx, 1–4.

Luiselli, L., Hema, E.M., Segniagbeto, G.H., Ouattara, V., Eniang, E.A., Di Vittorio, M., Amadi, N., Parfait, G., Pacini, N., Akani, G.C., Sirima, D., Guenda, W., Fakae, B.B., Dendi, D., Fa, J.E. (2019) Understanding the influence of non-wealth factors in determining bushmeat consumption: Results from four West African countries. Acta Oecologica, 94, 47–56.

Maritz, B., Alexander, G.J., Maritz, R.A. (2019) The underappreciated extent of cannibalism and ophiophagy in African cobras. Ecology, 100, e02522.

Medawar P.B. (1952) An Unsolved Problem in Biology. H.K. Lewis, London, UK.

Monaghan, P., Charmantier, A., Nussey, D. H., & Ricklefs, R. E. (2008). The evolutionary ecology of senescence. Functional Ecology, 22, 371–378.

Nussey, D. H., Froy, H., Lemaitre, J. F., Gaillard, J. M., & Austad, S. N. (2013). Senescence in natural populations of animals: widespread evidence and its implications for bio-gerontology. Ageing Research Reviews, 12, 214–225.

Oli, M. K. (2004). The fast–slow continuum and mammalian life-history patterns: an empirical evaluation. Basic and Applied Ecology, 5, 449–463.

Pradel, R. (1996). Utilization of capture-mark-recapture for the study of recruitment and population growth rate. Biometrics, 703–709.

Robert, K. A., & Bronikowski, A. M. (2010). Evolution of senescence in nature: physiological evolution in populations of garter snake with divergent life histories. The American Naturalist, 175, 147–159.

Royall, R. M. (1986). Model robust confidence intervals using maximum likelihood estimators. International Statistical Review/Revue Internationale de Statistique, 54, 221–226.

Salguero-Gómez, R., Jones, O. R. (2017). Life history trade-offs modulate the speed of senescence. In: Shefferson, R. P., Jones, O. R., Salguero-Gómez, R. (Eds.) The evolution of senescence in the tree of life. Pp 403–419. Cambridge University Press, UK.

Shefferson, R. P., Jones, O. R., & Salguero-Gómez, R. (Eds.). (2017). The evolution of senescence in the tree of life. Cambridge University Press, UK.

Stearns, S. C. (1992). The evolution of life histories. Oxford University Press, UK.

Tuberville, T. D., Norton, T. M., Todd, B. D., & Spratt, J. S. (2008). Long-term apparent survival of translocated gopher tortoises: a comparison of newly released and previously established animals. Biological Conservation, 141, 2690–2697.

Vaupel, J. W., Baudisch, A., Dölling, M., Roach, D. A., & Gampe, J. (2004). The case for negative senescence. Theoretical Population Biology, 65, 339–351.

Warner, D. A., Miller, D. A., Bronikowski, A. M., & Janzen, F. J. (2016). Decades of field data reveal that turtles senesce in the wild. Proceedings of the National Academy of Sciences, 113, 6502–6507.

Williams, G. C. (1957). Pleiotropy, natural selection, and the evolution of senescence. Evolution, 11, 398–411.

